## Supplementary Data for "The selfish yeast plasmid exploits a SWI/SNF-type chromatin remodeling complex for hitchhiking on chromosomes and ensuring high-fidelity propagation"

**Table S1. Yeast strains and their relevant features.** The genotypes of the yeast strains used in the present study along with the figures depicting the experimental results obtained with them are listed. Strains containing or lacking the native 2-micron plasmid are indicated as [Cir$^+$] or [Cir$^0$], respectively.

| Strains | Genotype/salient features | Relevant Figures |
|---|---|---|
| CMY685 | *MATa ade2-1 his3-11::*P$_{GAL10}$-*REP1*-P$_{GAL1}$-*REP2::HIS3 leu2-3,112 trp1 ura3-1* [Cir$^0$] | Figure 1, Figure S1, Figure 2 Figure S3 |
| CMY687 | *MATa ade2-1 his3-11::*P$_{GAL10}$-*3HA-REP1*-P$_{GAL1}$-*REP2-10Myc::HIS3 leu2-3,112 trp1 ura3-1* [Cir$^0$] | Figure 1, Figure S1, Figure 2 Figure S3 |
| CMY689 | *MATa ade2-1 his3-11::*P$_{GAL1}$-*REP1-10Myc*-P$_{GAL10}$-*3HA-REP2::HIS3 leu2-3,112 trp1 ura3-1* [Cir$^0$] | Figure 1, Figure S1, Figure 2 Figure S3 |
| CMY699 | *MATa ade2-1 his3-11::*P$_{GAL1}$-*FLP(Y343F)-10Myc::HIS3 leu2-3,112 trp1::*4x (half-*FRT*) *ura3-1* [Cir$^0$] | Figure 1, Figure S1, Figure 2 Figure S3 |
| CMY781 | *MATa pADE2 ade2-1 his3-11 leu2-3,112 trp1 ura3-1* [Cir$^0$] | Figure S4A |
| CMY782 | *MATa pADE2 ade2-1 his3-11 leu2-3,112 trp1 ura3-1 rsc2Δ::URA3* [Cir$^0$] | Figure S4A |
| CMY783 | *MATa pADE2 ade2-1 his3-11 leu2-3,112 trp1 ura3-1 truncated rsc2* [Cir$^0$] | Figure S4A |
| CMY795 | *MATa ade2-1 his3-11::*P$_{GAL10}$-*3HA-REP1*-P$_{GAL1}$-*REP2-10Myc::HIS3 leu2-3,112 trp1 ura3-1 rsc2Δ::URA3* [Cir$^0$] | Figure 4 Figure S4B-D, Figure 5, Figure6, Figure S6 |
| CMY797 | *MATa ade2-1 his3-11::*P$_{GAL10}$-*3HA-REP1*-P$_{GAL1}$-*REP2-10Myc::HIS3 leu2-3,112 trp1 ura3-1 truncated rsc2* [Cir$^0$] | Figure 4 Figure S4B-D, Figure 5, Figure 6, Figure S6 |
| CMY799 | *MATa ade2-1 his3-11::*P$_{GAL1}$-*REP1-10Myc*-P$_{GAL10}$-*3HA-REP2::HIS3 leu2-3,112 trp1 ura3-1 rsc2Δ::URA3* [Cir$^0$] | Figure 4 Figure S4B-D, Figure 5, Figure 6, Figure S6 |

| | | |
|---|---|---|
| CMY801 | *MATa ade2-1 his3-11::*P$_{GAL1}$*-REP1-10Myc-*P$_{GAL10}$*-3HA-REP2::HIS3 leu2-3,112 trp1 ura3-1 truncated rsc2* [Cir$^0$] | Figure 4 Figure S4B-D, Figure 5, Figure 6, Figure S6 |
| Y2HGold | *MATa trp1-901 leu2-3,112 ura3-52 his3-200 gal4Δ gal80Δ LYS2::GAL1*$_{UAS}$*–Gal1*$_{TATA}$*–His3 GAL2*$_{UAS}$*–Gal2*$_{TATA}$*–Ade2 URA3::MEL1*$_{UAS}$*–Mel1*$_{TATA}$ *AUR1-C MEL1* [Cir$^+$] | Figure 7, Figure S7–1 |
| Y187 | *MATα ura3-52 his3-200 ade2-101 trp1-901 leu2-3,112 gal4Δ gal80Δ met– URA3::GAL1*$_{UAS}$*–Gal1*$_{TATA}$*–LacZ, MEL1* [Cir$^+$] | Figure 7, Figure S7–1 |
| CMY891 | *MATa trp1-901 leu2-3,112 ura3-52 his3-200 gal4Δ gal80Δ LYS2::GAL1*$_{UAS}$*–Gal1*$_{TATA}$*–His3 GAL2*$_{UAS}$*–Gal2*$_{TATA}$*–Ade2 URA3::MEL1*$_{UAS}$*–Mel1*$_{TATA}$ *AUR1-C MEL1* [Cir$^0$] | Figure 7, Figure S7–1 |
| CMY893 | *MATα ura3-52 his3-200 ade2-101 trp1-901 leu2-3,112 gal4Δ gal80Δ met– URA3::GAL1*$_{UAS}$*–Gal1*$_{TATA}$*–LacZ MEL1* [Cir$^0$] | Figure 7, Figure S7–1 |
| CMY861 | *MATα ura3-52::*P$_{GAL10}$*-3HA-REP1-*P$_{GAL1}$*-REP2-10Myc::URA3 trp1 leu2Δ1 his3Δ200 pep4::HIS3 prb1Δ1.6R can1 RSC2-SGGGG-CBP-TEV-ProtA-ProtA::TRP1* [Cir$^+$] | Figure S7–5 |
| CMY859 | *MATα ura3-52::*P$_{GAL10}$*-3HA-REP1-*P$_{GAL1}$*-REP2-10Myc::URA3 trp1 leu2Δ1 his3Δ200 pep4::HIS3 prb1Δ1.6R can1 RSC1-SGGGG-CBP-TEV-ProtA-ProtA* [Cir$^+$] | Figure S7–7 |
| NPY7 | *MATα ura3-52::*P$_{GAL10}$*-3HA-REP1-*P$_{GAL1}$*-REP2-10Myc::URA3 trp1 leu2Δ1 his3Δ200 pep4::HIS3 prb1Δ1.6R can1 RSC2-SGGGG-CBP::TRP1* [Cir$^+$] | Figure S7–6 |
| NPY9 | *MATα ura3-52::*P$_{GAL10}$*-3HA-REP1-*P$_{GAL1}$*-REP2-10Myc::URA3 trp1 leu2Δ1 his3Δ200 pep4::HIS3 prb1Δ1.6R can1 RSC1-SGGGG-CBP::TRP1* [Cir$^+$] | Figure S7–8 |
| SGY110 81 | *MATa/α ade2-1 trp1-1 can1-100 leu2-3,112 his3-11 ura3-1* [Cir$^+$] | Figure 8, Figures S8–1-4 |
| SGY110 82 | *MATa*/α *ade2-1 trp1-1 can1-100 leu2-3,112 ura3-1 RSC2-6HA::URA3/RSC2-6HA::URA3 his3-11::P*$_{GAL1}$*-REP1-10Myc::HIS3* [Cir$^+$] | Figure 8, Figure S8–1 |
| SGY110 83 | *MATa*/α *ade2-1 trp1-1 can1-100 leu2-3,112 ura3-1 RSC2-6HA::URA3/RSC2-6HA::URA3 his3-11::*P$_{GAL1}$*-REP1-10Myc::HIS3* [Cir$^0$] | Figure S8–3 |
| SGY110 84 | *MATa*/α *ade2-1 trp1-1 can1-100 leu2-3,112 ura3-1 RSC2-6HA::URA3/RSC2-6HA::URA3 his3-11::P*$_{GAL1}$*-REP2-10Myc::HIS3* [Cir$^+$] | Figure 8, Figure S8–1 |
| SGY110 85 | *MATa*/α *ade2-1 trp1-1 can1-100 leu2-3,112 ura3-1 RSC2-6HA::URA3/RSC2-6HA::URA3 his3-11::*P$_{GAL1}$*-REP2-10Myc::HIS3* [Cir$^0$] | Figure S8–3 |

| | | |
|---|---|---|
| SGY11086 | *MATa*/α *ade2-1 trp1-1 can1-100 leu2-3,112 ura3-1 RSC1-6HA::KanMX4/RSC1-6HA::KanMX4 his3-11::*P$_{GAL1}$*-REP1-10Myc::HIS3* [Cir$^{+}$] | Figure 8, Figure S8–1 |
| SGY11087 | *MATa*/α *ade2-1 trp1-1 can1-100 leu2-3,112 ura3-1 RSC1-6HA::KanMX4/RSC1-6HA::KanMX4 his3-11::*P$_{GAL1}$*-REP1-10Myc::HIS3* [Cir$^{0}$] | Figure S8–3 |
| SGY11088 | *MATa*/α *ade2-1 trp1-1 can1-100 leu2-3,112 ura3-1 RSC1-6HA::KanMX4/RSC1-6HA::KanMX4 his3-11::*P$_{GAL1}$*-REP2-10Myc::HIS3* [Cir$^{+}$] | Figure 8, Figure S8–1 |
| SGY11089 | *MATa*/α *ade2-1 trp1-1 can1-100 leu2-3,112 ura3-1 RSC1-6HA::KanMX4/RSC1-6HA::KanMX4 his3-11::*P$_{GAL1}$*-REP2-10Myc::HIS3* [Cir$^{0}$] | Figure S8–3 |
| SGY11090 | *MATa*/α *ade2-1 trp1-1 can1-100 leu2-3,112 ura3-1 his3-11 RSC2-6HA::URA3 RSC1-9Myc::KanMX4* [Cir$^{+}$] | Figure S8–2 |
| SGY11091 | *MATa*/α *trp1-1 can1-100 leu2-3,112 his3-11 ura3-1 RSC2-9Myc::KanMX4/RSC2-9Myc::KanMX4 ade2-1::GFP-LacI::ADE2/ade2-1::GFP-LacI::ADE2* [Cir$^{+}$] | Figure 8, Figure S8–1 |
| SGY11092 | *MATa*/α *trp1-1 can1-100 leu2-3,112 his3-11 ura3-1 RSC2-9Myc::KanMX4/RSC2-9Myc::KanMX4 ade2-1::GFP-LacI::ADE2/ade2-1::GFP-LacI::ADE2* [Cir$^{0}$] | Figure S8–3 |
| SGY11093 | *MATa*/α *trp1-1 can1-100 leu2-3,112 his3-11 ura3-1 RSC1-9Myc::KanMX4/RSC1-9Myc::KanMX4 ade2-1::GFP-LacI:: ADE2/ade2-1::GFP-LacI::ADE2* [Cir$^{+}$] | Figure 8, Figure S8–1 |
| SGY11094 | *MATa*/α *trp1-1 can1-100 leu2-3,112 his3-11 ura3-1 RSC1-9Myc::KanMX4/RSC1-9Myc::KanMX4 ade2-1::GFP-LacI:: ADE2/ade2-1::GFP-LacI::ADE2* [Cir$^{0}$] | Figure S8–3 |
| SGY11095 | *MATa*/α *can1-100 leu2-3,112 trp1-1* P$_{GAL1}$*-NDT80::TRP1* P$_{GDP1}$*-GAL4-ER RSC2-6HA::KanMX4 his3-11::* P$_{GAL1}$*-REP1-10Myc::HIS3 ura3-1::*P$_{GAL1}$*-REP2::URA3* [Cir$^{0}$] | Figure S8–4 |
| SGY11096 | *MATa*/α *can1-100 leu2-3,112 trp1-1* P$_{GAL1}$*-NDT80::TRP1* P$_{GPD1}$*-Gal4-ER RSC1-6HA::KanMX4 his3-11::*P$_{GAL1}$*-REP2-10Myc::HIS3 ura3-1::*P$_{GAL1}$*-REP1::URA3* [Cir$^{0}$] | Figure S8–4 |
| SGY11099 | *MATa trp1-1 can1-100 his3-11 tRNA-Val::gRNA-TetO leu2-3,112::TetR-tdTomato::LEU2 ade2-1::GFP-LacI-ADE2 ura3::P$_{GAL10}$-REP1-P$_{GAL1}$-REP2::URA3* [Cir$^{0}$] | Figure 9 |
| SGY11100 | *MATa trp1-1 can1-100 his3-11 ura3-1 tRNA-Val::gRNA-TetO leu2-3,112::TetR-tdTomato::LEU2 ade2-1::GFP-LacI-ADE2* [Cir$^{0}$] | Figure 9 |
| SGY11097 | *MATa trp1-1 can1-100 leu2-3,112 his3-11 ura3-1 ade2-1::GFP-LacI::ADE2* [Cir$^{0}$] | Figure 10 |
| SGY11098 | *MATa trp1-1 can1-100 leu2-3,112 his3-11 ura3-1 ade2-1::GFP-LacI::ADE2 SFH1::SFH1-LacI* [Cir$^{0}$] | Figure 10 |

**Table S2. Plasmids used in this study.** The plasmids listed below as groups I-III were used for yeast two-hybrid analyses, expression in *E. coli* and interaction assays, and cell biological/fluorescence microscopy experiments in yeast, respectively. The figures containing experimental results to which they contributed are indicated. For previously described plasmids, the relevant references are given.

| Plasmids | Relevant Figures |
|---|---|
| **Group I** | |
| pGBKT7 (DNA binding domain vector in two-hybrid assays) | Figure 7, Figure S7–1 |
| pGADT7 (activation domain vector in two-hybrid assays) | Figure 7, Figure S7–1 |
| pGBKT7-*REP1* | Figure 7, Figure S7–1 |
| pGADT7-AD-*REP1* | Figure 7, Figure S7–1 |
| pGBKT7-*REP2* | Figure 7, Figure S7–1 |
| pGADT7-AD-*REP2* | Figure 7, Figure S7–1 |
| pGBKT7-*SFH1* | Figure 7, Figure S7–1 |
| pGADT7-*SFH1* | Figure 7, Figure S7–1 |
| pGBKT7-Sfh1(Δ1-278) | Figure 7, Figure S7–1 |
| pGADT7-Sfh1(Δ1-278) | Figure 7, Figure S7–1 |
| **Group II** | |
| pET28a-His6-*SFH1* | Figure S7–2 |

| | |
|---|---|
| pET28a-His6-Sfh1(Δ1-278) | Figure S7–3 |
| pGEX2T-*REP1* | Figures S7–2 and 3 |
| pGEX2T-*REP2* | Figures S7–2 and 3 |
| pGEX2T-GST-*REP2*- reverse P-tac-3HA-*REP1* | Figure S7–4 |
| **Group III** | |
| pSV1-*ORI-STB* [LacO]$_{256}$ (*LEU2*) | Figures S8–1 and 3 (Mehta *et al*. 2002 J Cell Biol 158: 625-637) |
| pBM272-*REP1* (*ARS-CEN* $P_{GAL10}$-*REP1*) | Figure S8–4 (Yang *et al*. 2004 Mol Cell Biol: 24: 5290-5303) |
| pBM272-*REP2* (*ARS-CEN* $P_{GAL10}$-*REP2*) | Figure S8–4 (Yang *et al*. 2004 Mol Cell Biol: 24: 5290-5303) |
| pSG1: $P_{GAL1}$-*CEN3-STB-ORI* cloned in pSV5) | Figures 9 and 10 (Ghosh *et al*. 2010 PNAS 104: 13034-13039; Mehta *et al*. 2002 J Cell Biol 158: 625-637) |
| pSV5: *ORI-STB* [LacO]$_{256}$ (*TRP1*) | Figure 10 (Mehta *et al*. 2002 J Cell Biol 158: 625-637) |

Figure S1

A

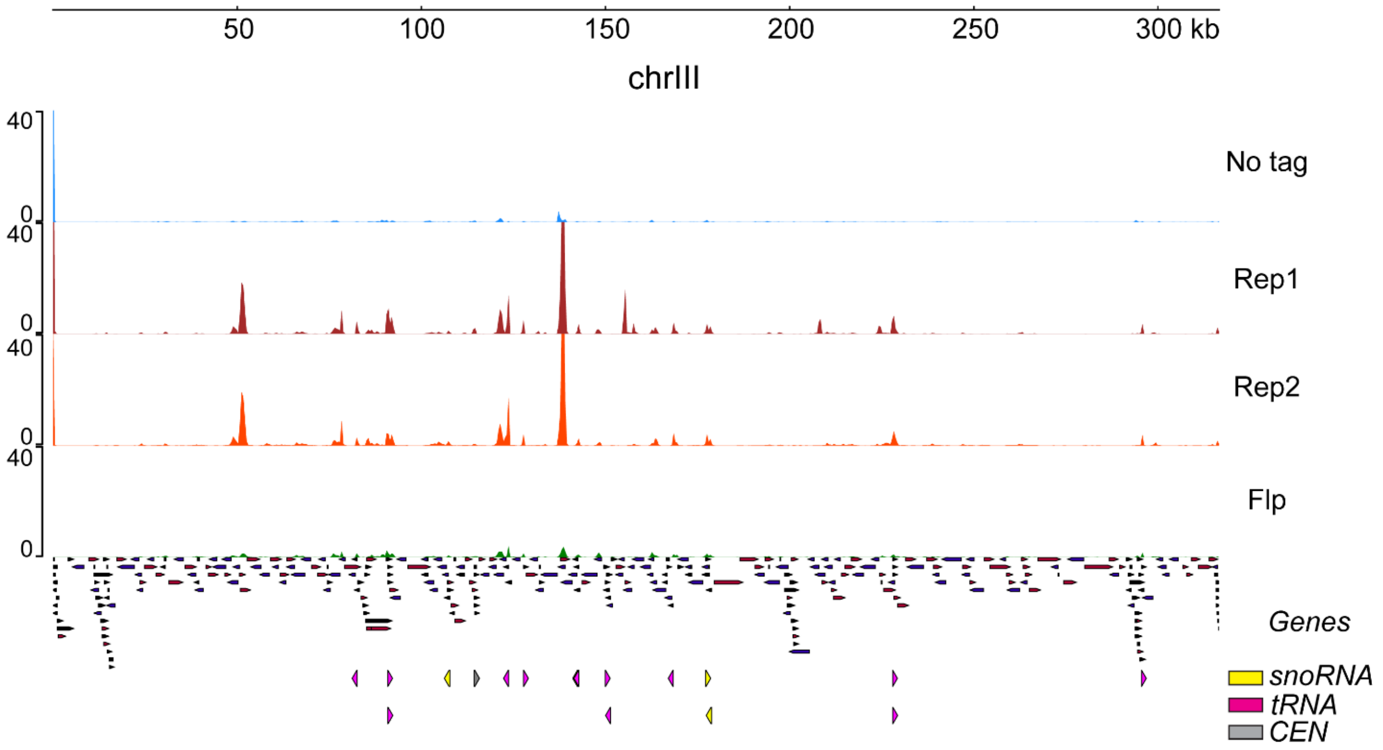
50
100
150
200
250
300 kb
chrIII
40
0
No tag
Rep1
Rep2
Flp
Genes
snoRNA
tRNA
CEN

B

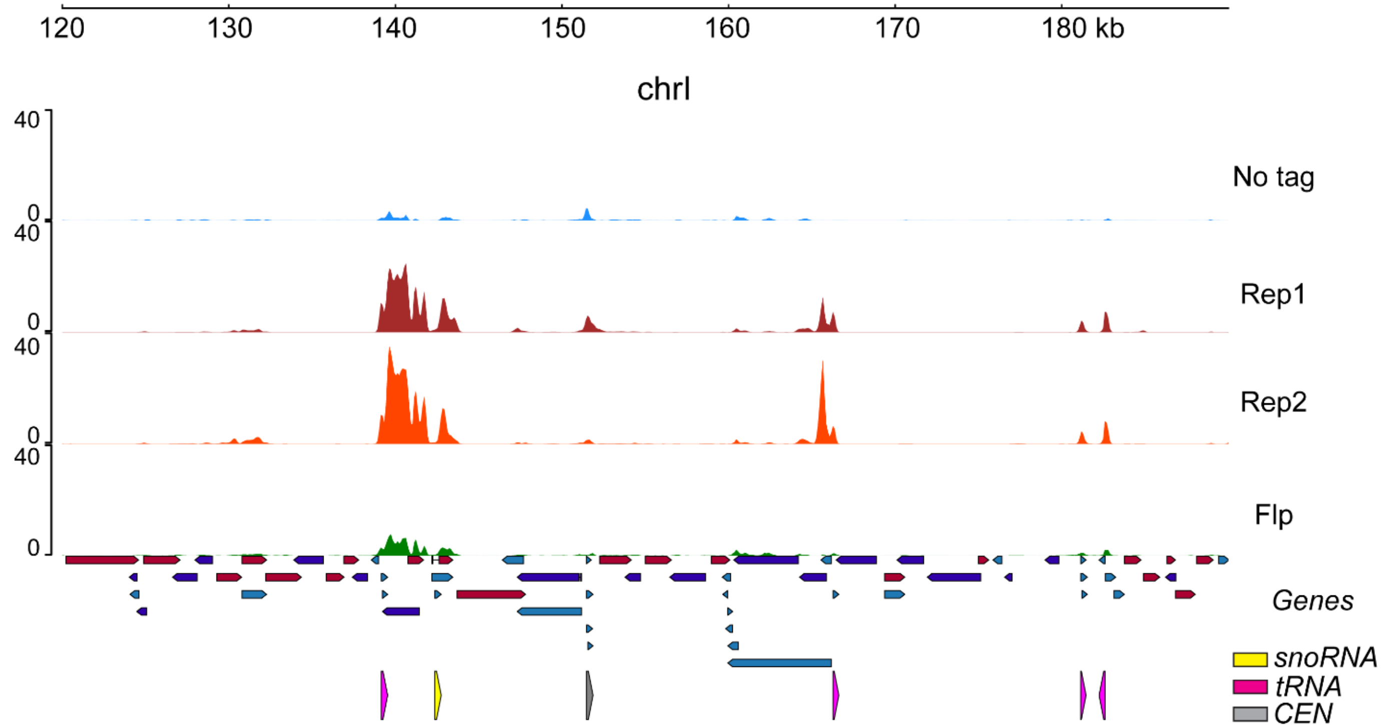
120
130
140
150
160
170
180 kb
chrI
40
0
No tag
Rep1
Rep2
Flp
Genes
snoRNA
tRNA
CEN

**Figure S1. Overview of Rep protein localization.** Binding of Rep1 and Rep2 assayed by ChIP-seq is shown using input-corrected signal tracks along with negative controls (No tag and Flp). The locations of snoRNAs, tRNAs and *CEN* are shown below each track using the colors indicated. **(A)** chrIII. **(B)** A region of chrI.

Figure S3

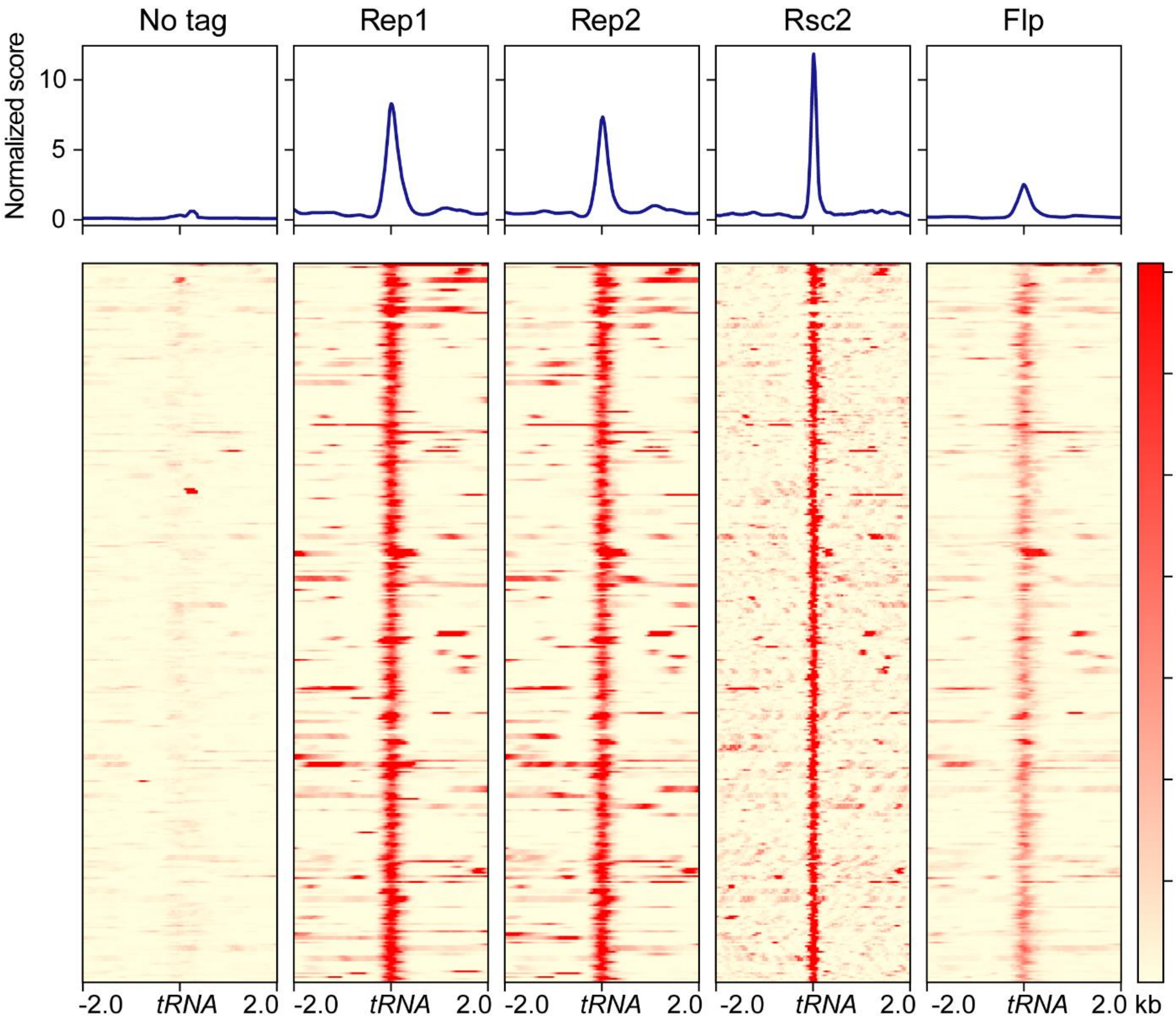

**Figure S3.** Rep protein and Rsc2 localization at tRNA loci. The top part of each panel shows the average binding profile and the bottom shows binding to each region as a heat map.

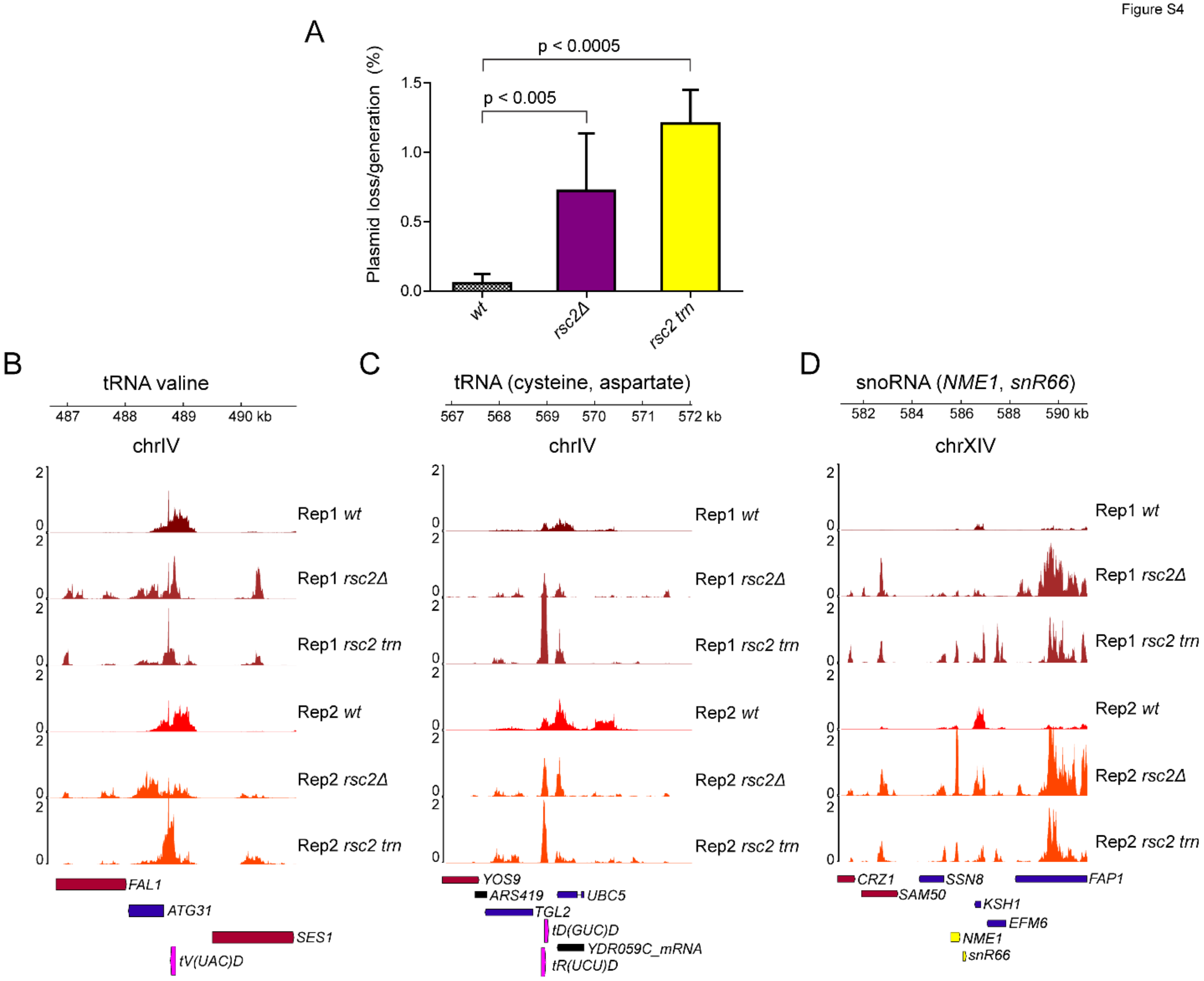

**Figure S4. (A)** Plasmid loss rate per generation, measured as described previously (Murray and Cesareni, 1986), is increased in both *rsc2Δ* and truncated rsc2 (rsc2 trn). **(B-D)** Rep1 and Rep2 localization at tRNA and snoRNA loci in wild-type, *rsc2Δ* and truncated rsc2 strains as measured by ChIP-seq. In some instances, a redistribution of Rep1 and Rep2 can be seen around the original binding sites in the rsc2 mutant strains.

Figure S6

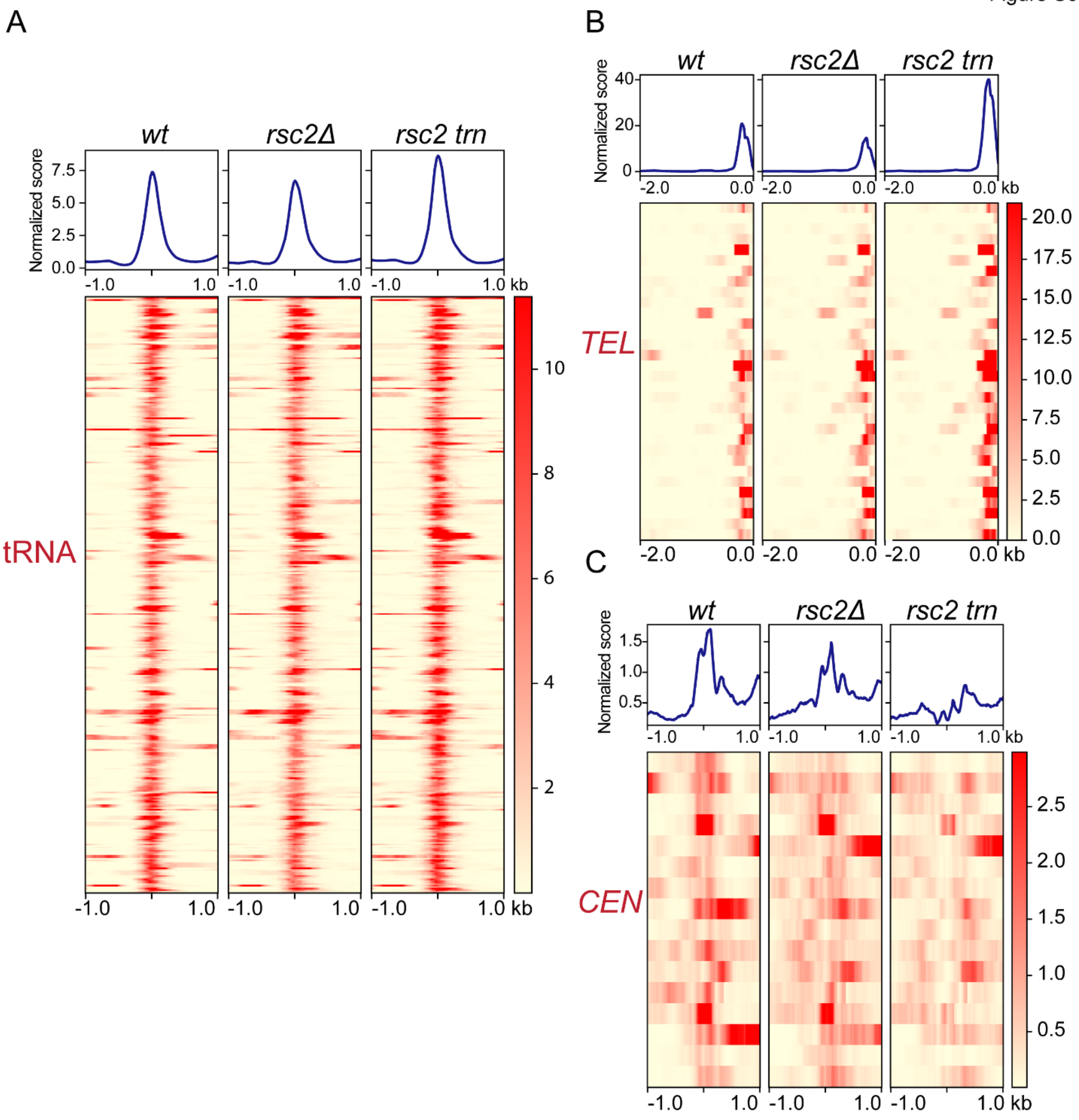

**Figure S6. Rep2 localization at tRNAs, *TEL*s and *CEN*s under *rsc2* mutation.** Binding of Rep1 at **(A)** tRNA loci, **(B)** *TEL*s and **(C)** *CEN*s as measured by ChIP-seq.

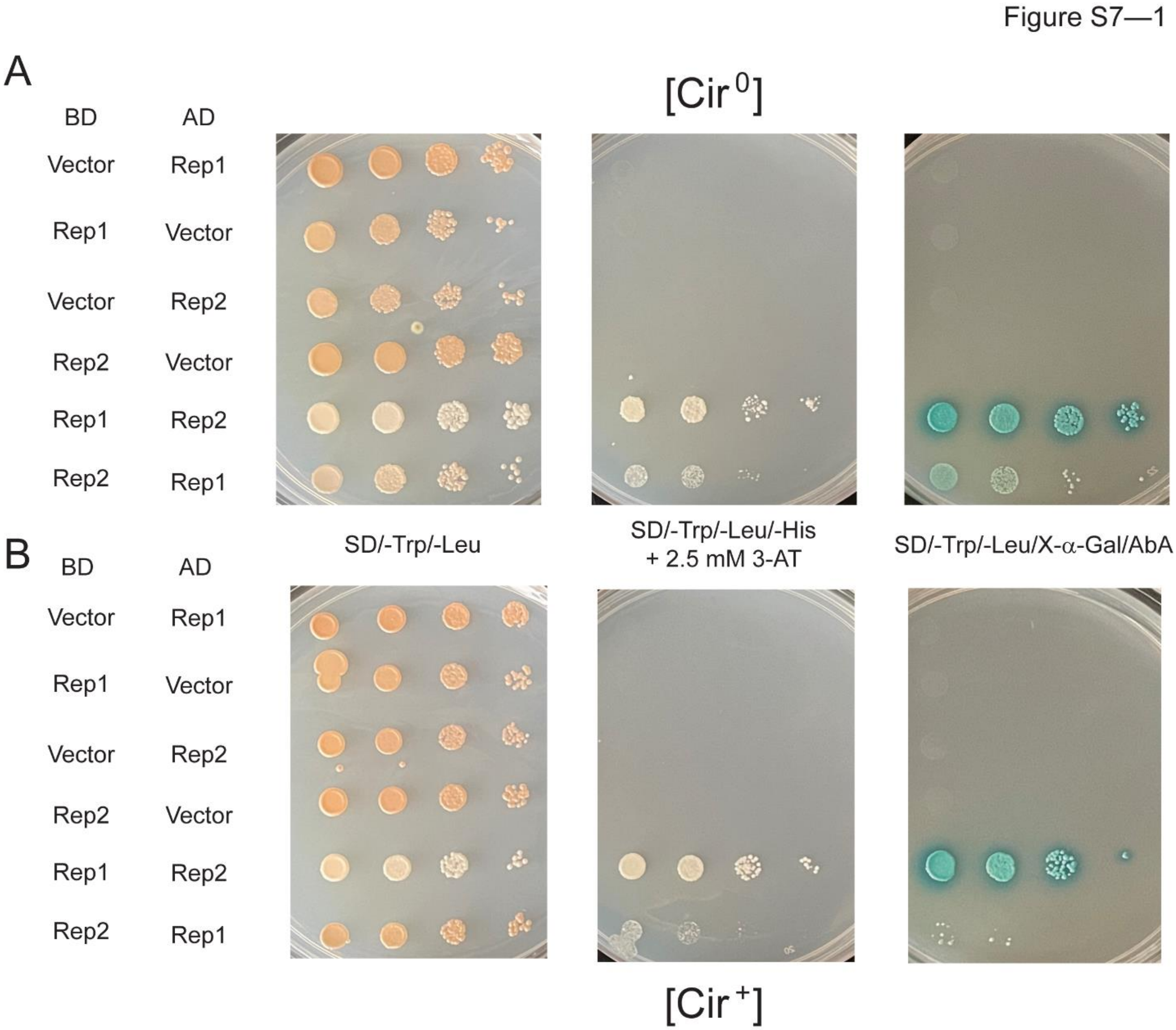

**Figure S7–1**. **Rep1-Rep2 interaction in the two-hybrid assay**. **(A, B)** The interaction between Rep1 and Rep2 tested here served as a positive control for the assays shown in Figure 7. This interaction was weaker with the [Rep2-BD]-[Rep1-AD] pair than with the [Rep1-BD]-[Rep2-AD] pair in both the [Cir$^0$] and [Cir$^+$] hosts. Presumably, the domain fusions in the former bait-prey configuration interferes partially with Rep1-Rep2 interaction.

Figure S7—2

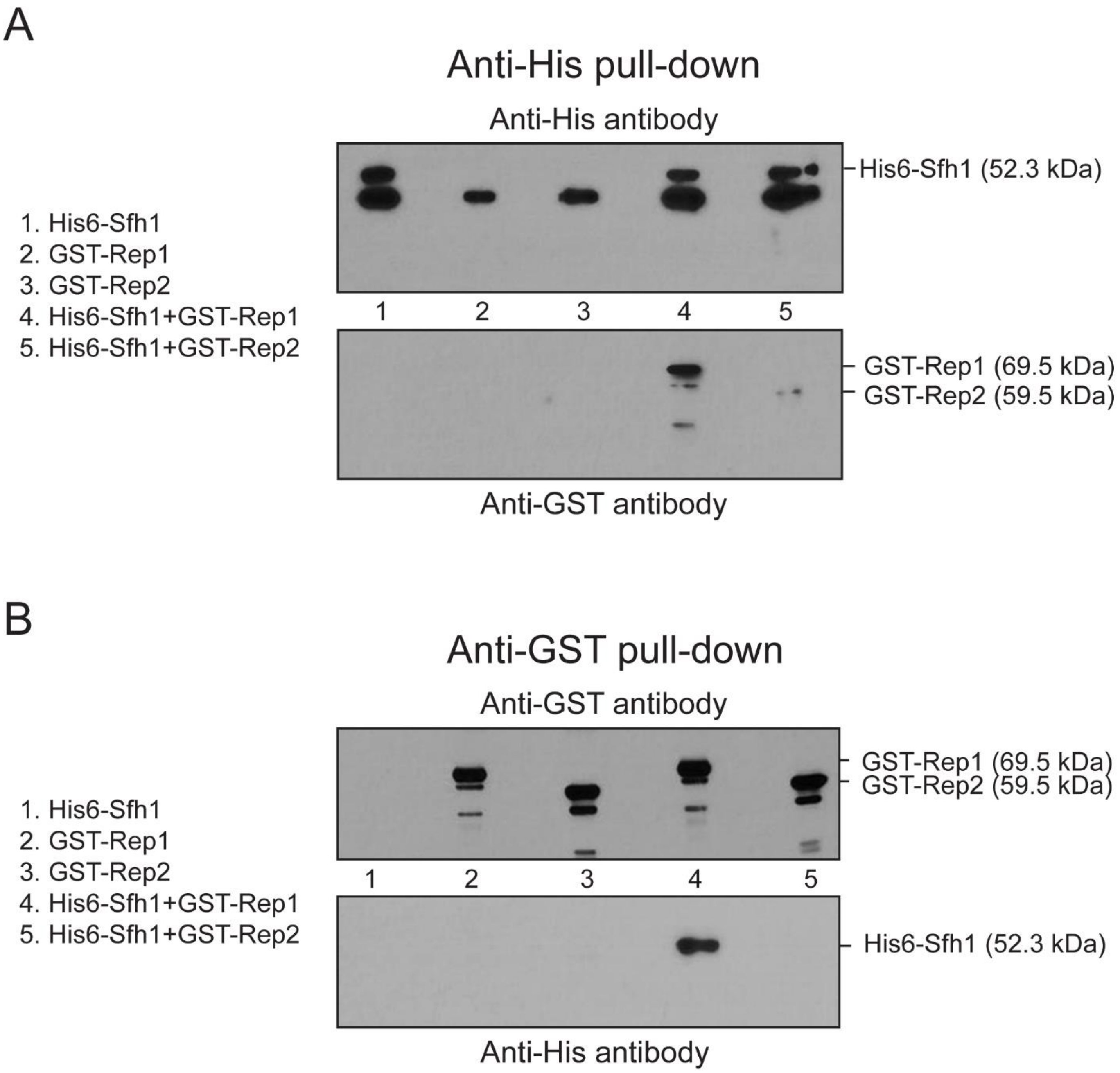

**Figure S7–2**. **Interaction of Sfh1 with Rep1 or Rep2 when these proteins are expressed in *E. coli*.** The indicated proteins were expressed in *E. coli* individually or in pairs, and were immunoprecipitated using Anti-His **(A)** or Anti-GST **(B)** antibodies. Co-immunoprecipitation of partner proteins was probed by western blot analysis.

Figure S7—3

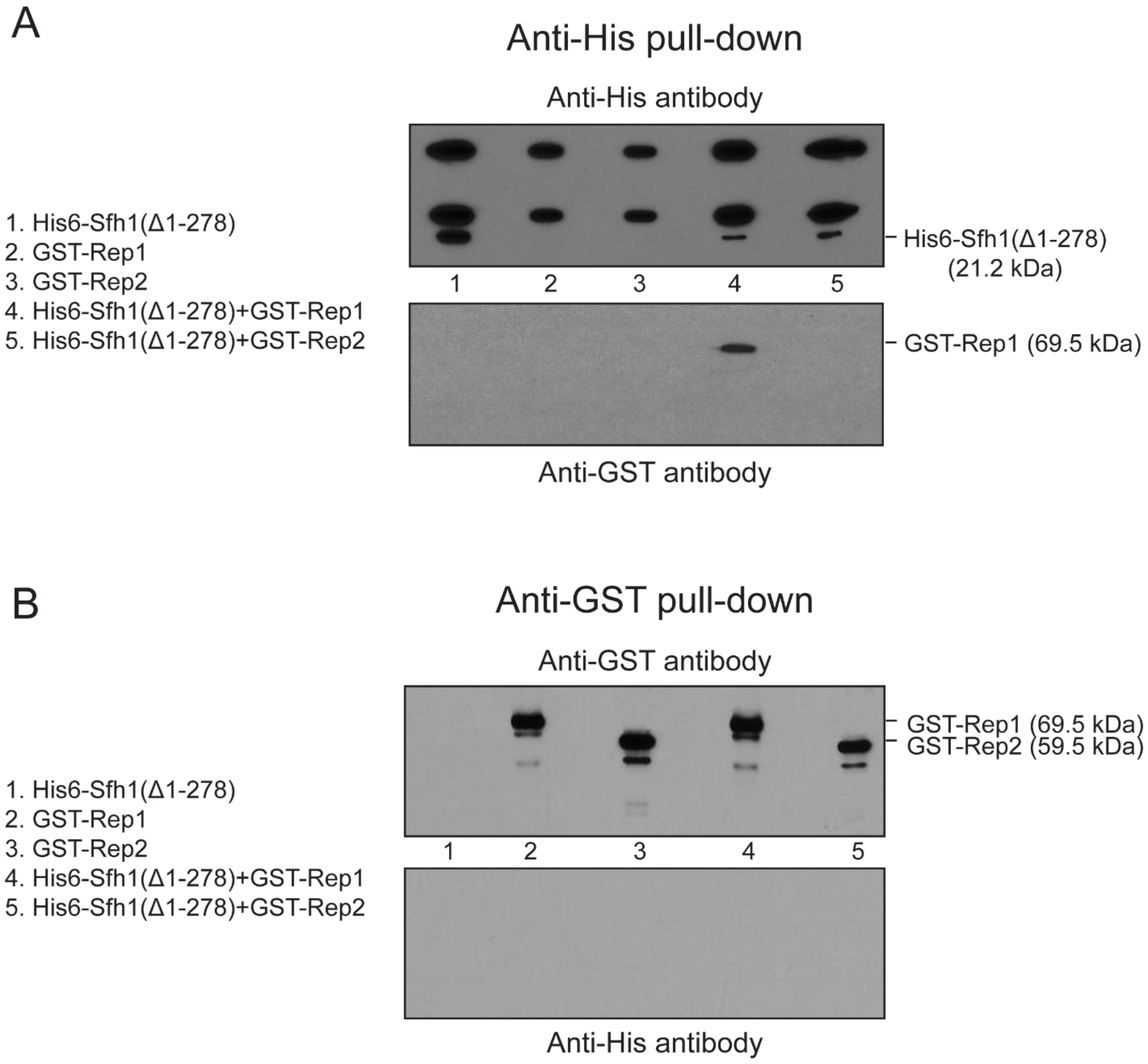

**Figure S7–3. Rep1-Sfh1(Δ1-278) and Rep2-Sfh1(Δ1-278) interactions assayed using *E. coli*-expressed proteins. (A, B)** The assays were similar to those shown in Figure S7–2, except that Sfh1(Δ1-278) was expressed instead of Sfh1.

Figure S7—4

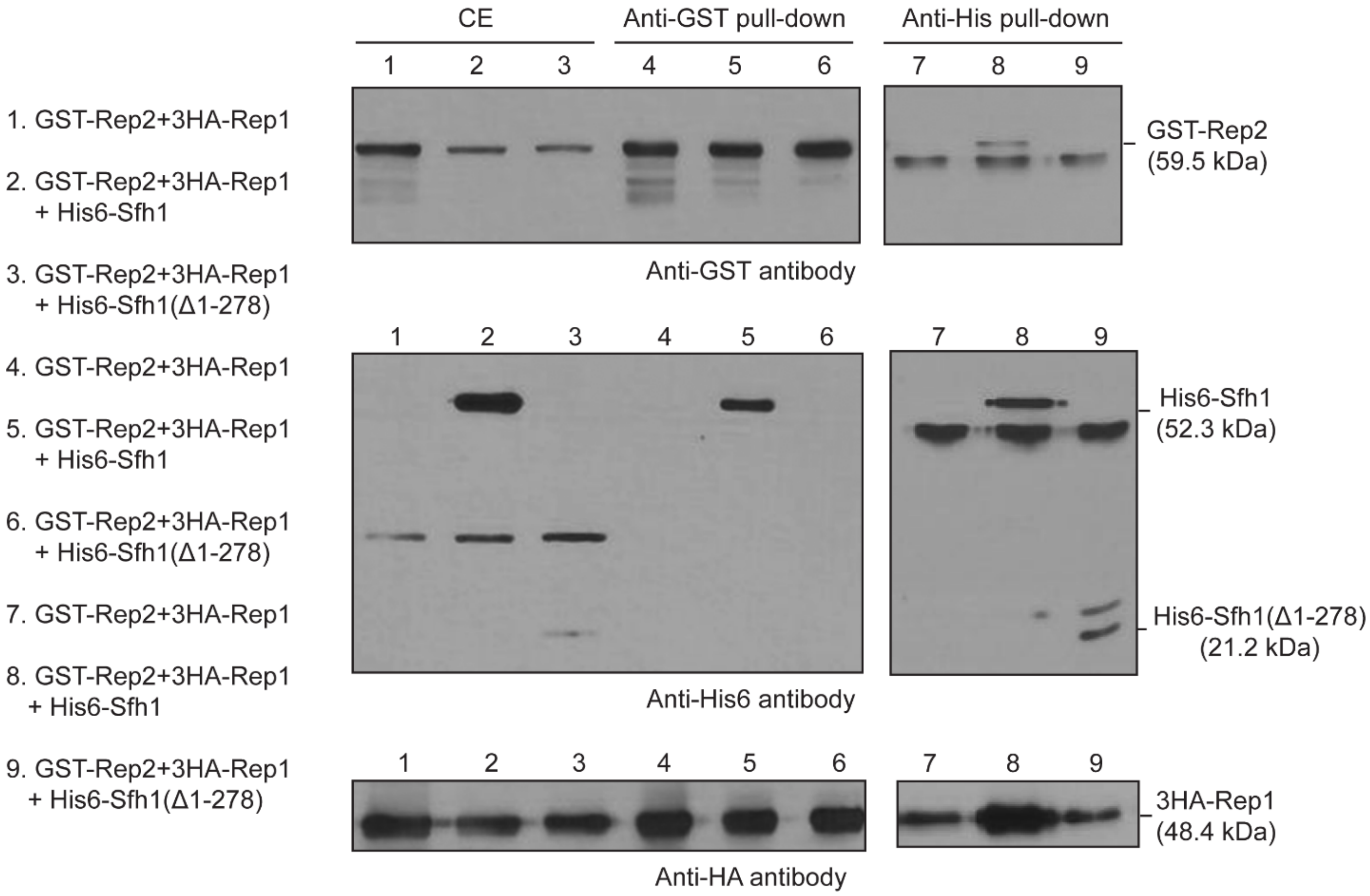

**Figure S7–4. Interactions of Sfh1 or Sfh1(Δ1-278) with Rep1 or Rep2 when the Rep proteins are co-expressed**. **(A, B)** The analyses were performed using extracts from *E. coli* strains expressing either Sfh1 or Sfh1(Δ1-278) together with both Rep1 and Rep2. The methodologies were analogous to those employed for the assays shown in Figures S7–2 and 3.

Figure S7—5

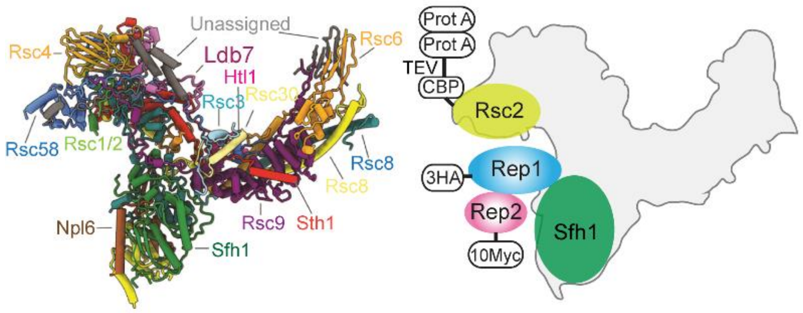

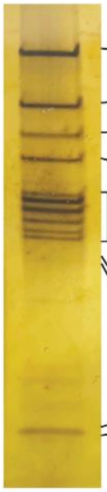

| Gene | Total peptides | Total Spectral Count | % |
|---|---|---|---|
| STH1 | 106 | 1699 | 73.4 |
| RSC1 | 5 | 23 | 1.0 |
| RSC2 | 83 | 1372 | 59.2 |
| RSC3 | 53 | 470 | 20.3 |
| RSC30 | 43 | 536 | 23.1 |
| RSC4 | 53 | 754 | 32.6 |
| RSC9 | 38 | 626 | 27.0 |
| RSC8 | 57 | 2316 | 100.0 |
| RSC58 | 50 | 621 | 26.8 |
| RSC6 | 39 | 775 | 33.5 |
| ARP7 | 45 | 796 | 34.4 |
| ARP9 | 36 | 590 | 25.5 |
| NPL6 | 34 | 559 | 24.1 |
| SFH1 | 23 | 797 | 34.4 |
| LDB7 | 8 | 251 | 10.8 |
| RTT102 | 10 | 118 | 5.1 |
| HTL1 | 9 | 69 | 3.0 |
| REP1 | 16 | 46 | 2.0 |
| REP2 | 7 | 17 | 0.7 |

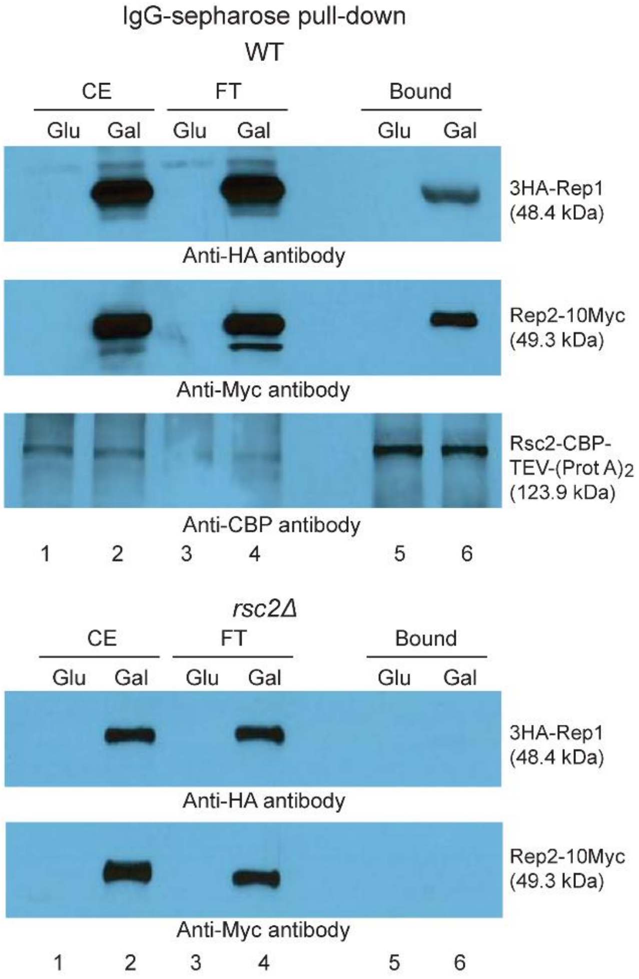

**Figure S7–5. Interactions of Rep1 or Rep2 with affinity-enriched RSC2 complex.** The cryo-EM structure of the yeast RSC complex (Patel et al., 2019) (RCSB PDB 6V8O) and an idealized rendition of the potential interaction of Rep1-Rep2 with the RSC2 complex are diagrammed at the top left. The epitope tags fused to Rep1 (HA), Rep2 (Myc) and Rsc2 are indicated. The CBP (calmodulin binding peptide)-tag on Rsc2 was separated from the two tandem Prot A (Protein A)-tags by the TEV protease cleavage sequence. A silver-stained profile of the RSC2 complex following affinity enrichment using

IgG-sepharose and fractionation by SDS-polyacrylamide gel electrophoresis is shown below at the left. The relevant mass spectrometry data are shown in the adjacent Table. Results from western blot probing of the enriched RSC2 complex from a wild type (WT) strain using anti-HA and anti-Myc antibodies (to target Rep1 and Rep2, respectively) are displayed at the right. As a negative control, mock enrichment was performed in an *rsc2Δ* strain. In the assays shown here and in Figures S7–6-8, the expression of the tagged Rep1 and Rep2 was controlled by the *GAL* promoter.

Figure S7—6

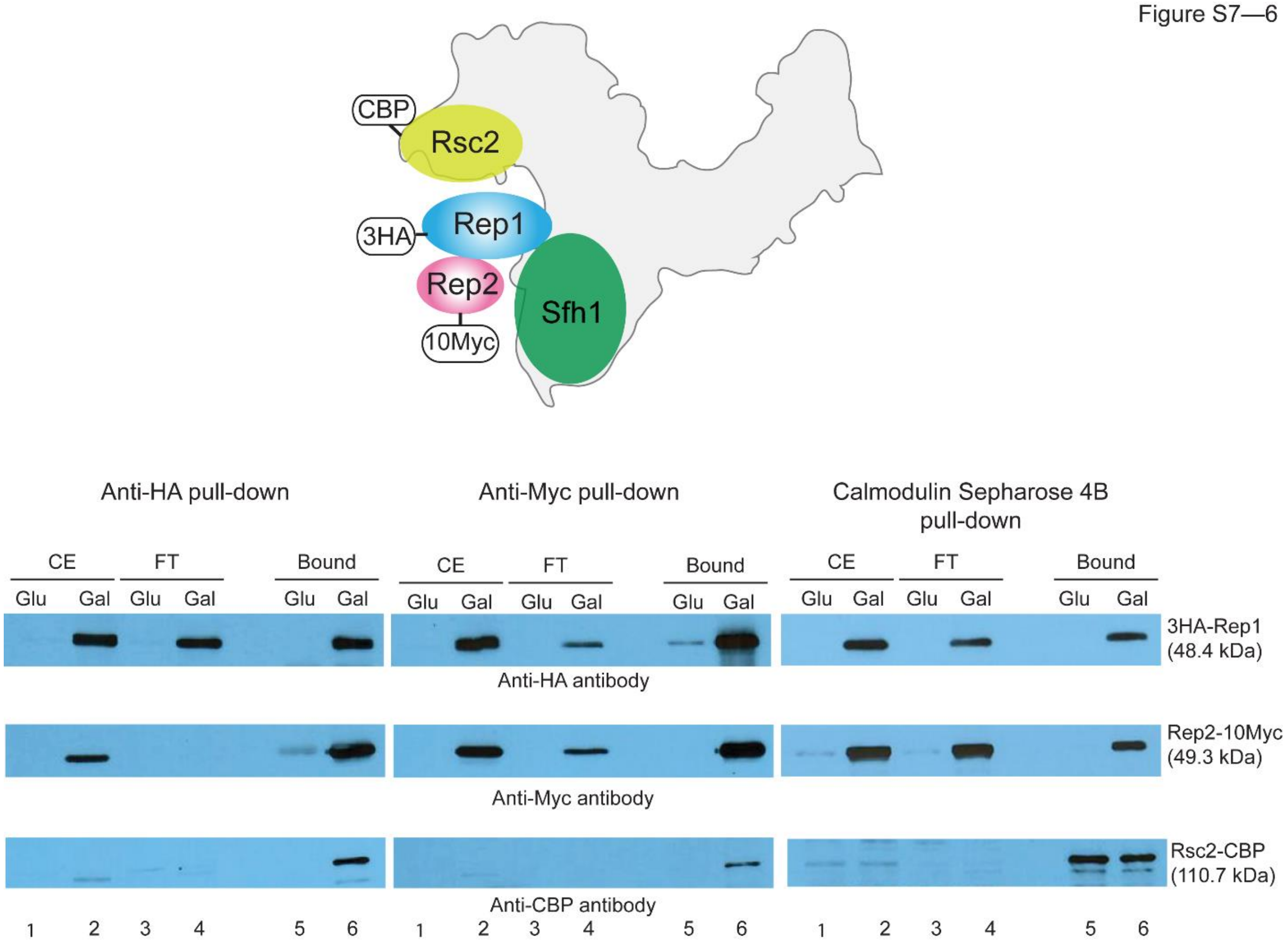

**Figure S7–6. Verification of [Rep1-Rep2]-RSC2 complex interaction by individual enrichment of Rep1, Rep2 or Rsc2.** The schematic diagram depicting the potential interaction of the RSC2 complex with Rep1-Rep2 is modeled after the corresponding diagram in Figure S7–5. The Rsc2 protein derivative for this assay carried the CBP-tag without the accompanying dual Protein A-tag. The primary enrichment was performed using anti-HA (left panel), anti-Myc (middle panel) or anti-CBP (right panel) antibody. Western blotting was performed with each of these antibodies to test the co-enrichment of suspected partner proteins.

Figure S7—7

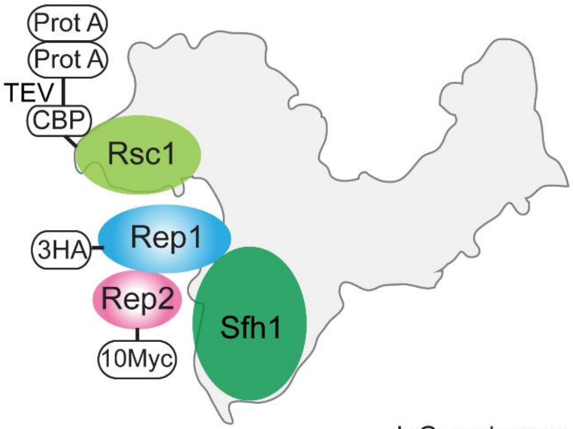

| Gene | Total peptides | Total Spectral Count | % |
|---|---|---|---|
| STH1 | 79 | 492 | 100.0 |
| RSC1 | 44 | 340 | 69.1 |
| RSC2 | 5 | 6 | 1.2 |
| RSC3 | 37 | 173 | 35.2 |
| RSC30 | 28 | 195 | 39.6 |
| RSC4 | 40 | 203 | 41.3 |
| RSC9 | 26 | 129 | 26.2 |
| RSC8 | 43 | 453 | 92.1 |
| RSC58 | 32 | 243 | 49.4 |
| RSC6 | 32 | 238 | 48.4 |
| ARP7 | 27 | 237 | 48.2 |
| ARP9 | 26 | 107 | 21.7 |
| NPL6 | 34 | 223 | 45.3 |
| SFH1 | 14 | 201 | 40.9 |
| LDB7 | 4 | 32 | 6.5 |
| RTT102 | 14 | 55 | 11.2 |
| HTL1 | 6 | 19 | 3.9 |
| REP1 | 13 | 34 | 6.9 |
| REP2 | 15 | 72 | 14.6 |

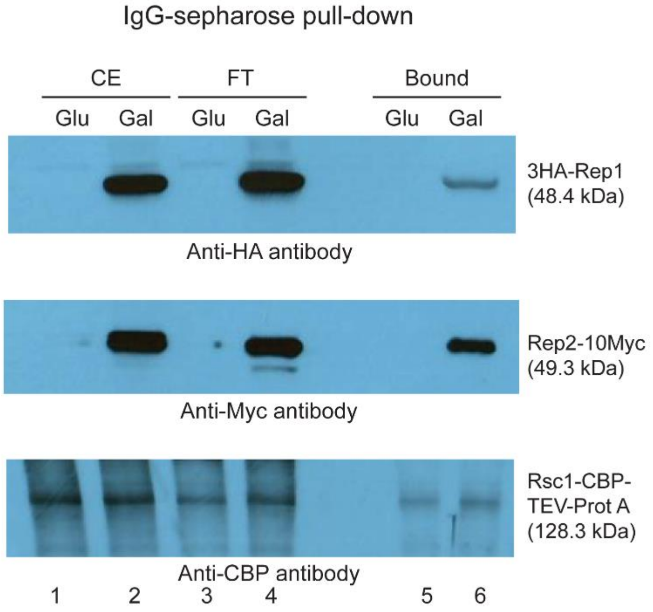

**Figure S7–7. Interactions of Rep1 or Rep2 with the affinity-enriched RSC1 complex.** The schematic diagram (top) for Rep1-Rep2 interaction with the RSC1 complex is a close replica of that in Figure S7–5, with Rsc2 replaced by Rsc1. The epitope tags fused to Rep1, Rep2 and Rsc1 are indicated. The Table below at the left lists the relevant mass spectrometry data for the RSC1 complex obtained by enrichment on IgG-sepharose beads. The association of Rep1 or Rep2 with the enriched complex was probed using anti-HA or anti-Myc antibodies (directed to Rep1 or to Rep2, respectively) (right panel).

Figure S7—8

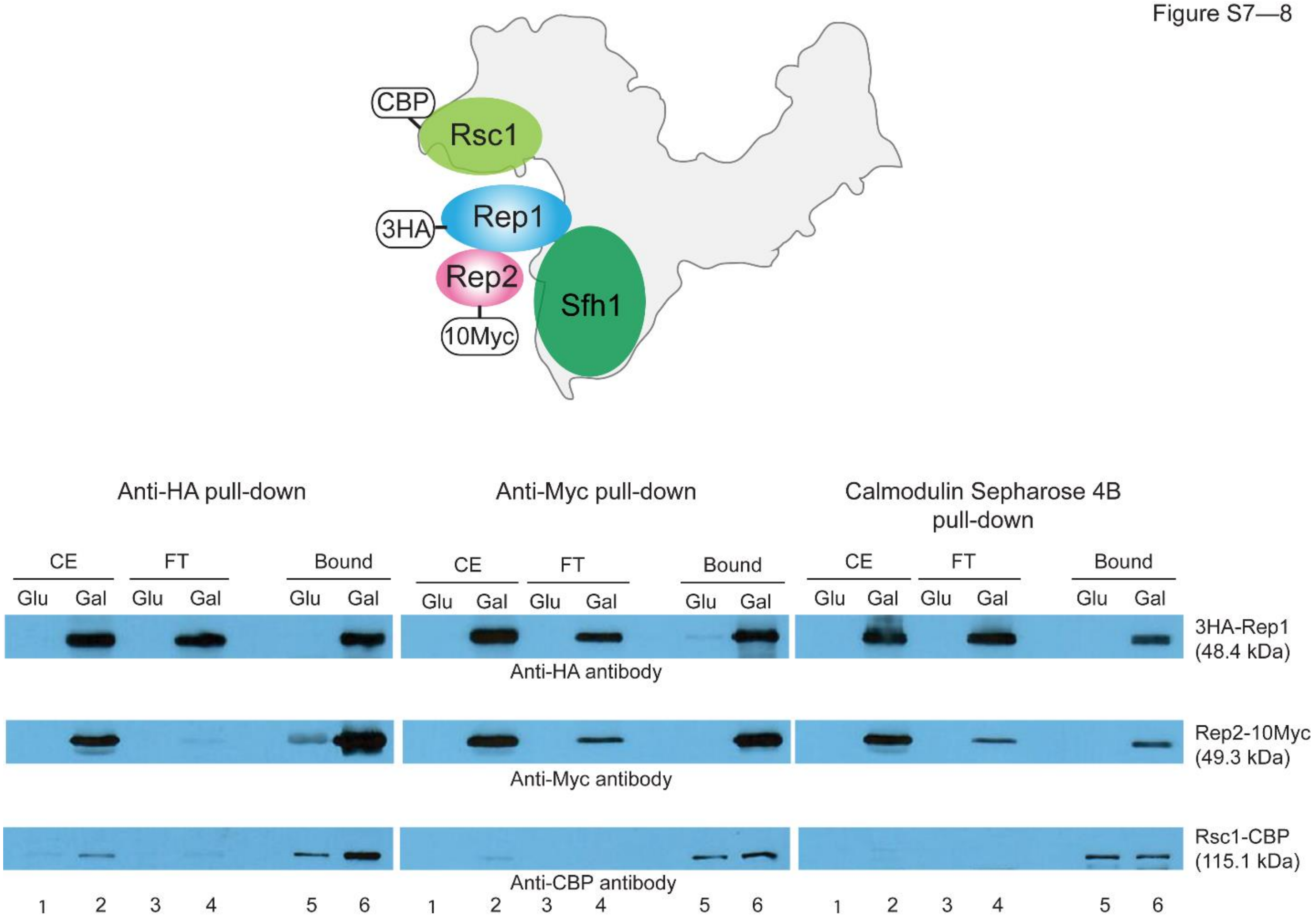

**Figure S7–8. Verification of [Rep1-Rep2]-RSC1 complex interaction by individual enrichment of Rep1, Rep2 or Rsc1.** The schematic diagram from Figure S7–6 is redrawn here with Rsc1-CBP replacing Rsc2-CBP. Enrichment of individual proteins and probing for associated proteins by western blotting were carried out as in the assays depicted in Figure S7–6.

Figure S8—1

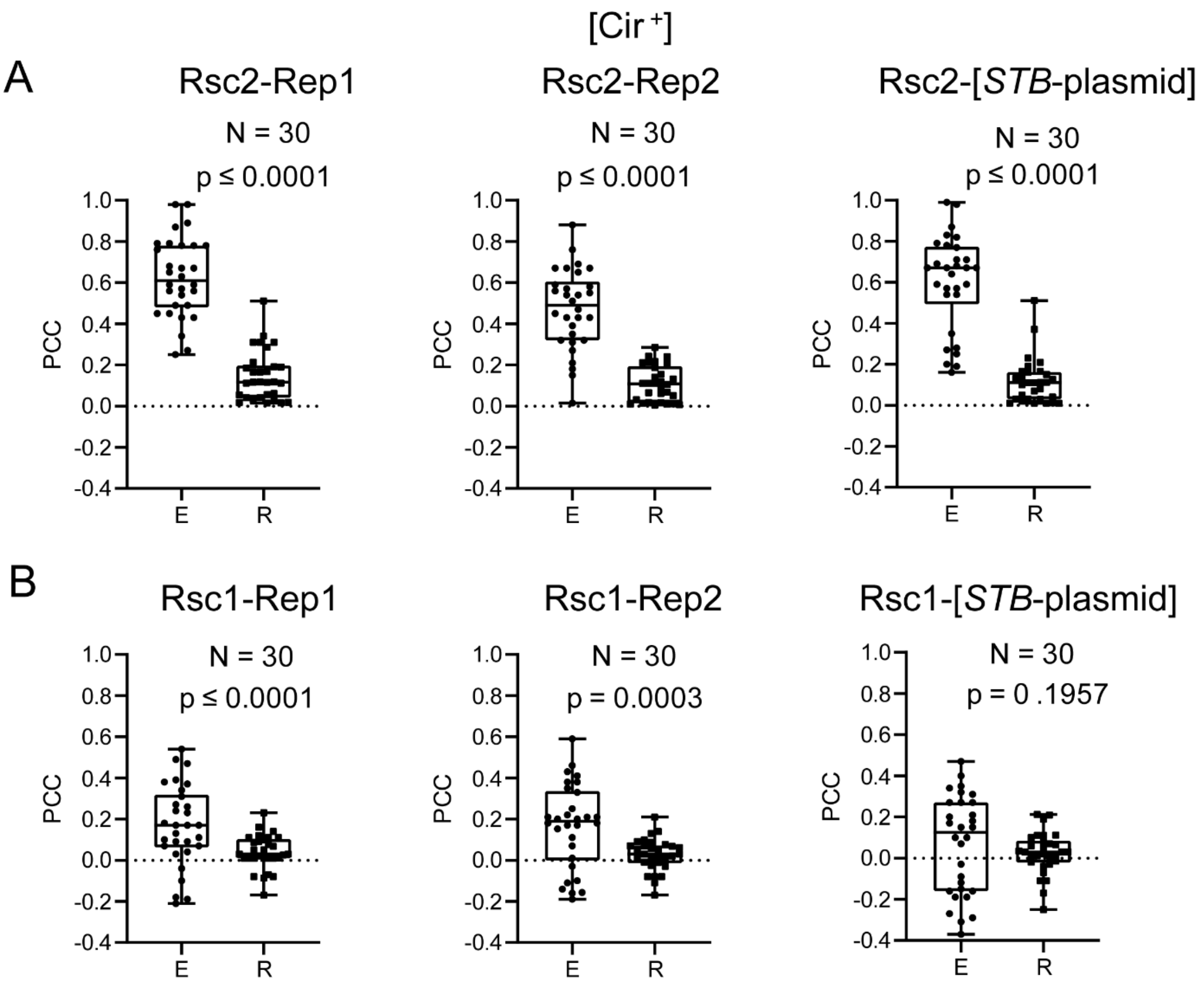

**Figure S8–1. Overlap between Rsc1 or Rsc2 fluorescence and Rep1 or Rep2 fluorescence in [Cir$^+$] chromosome spreads relative to fluorescence overlap by chance. (A, B)** The experimental protocols were as described under Figure 8. To obtain the degree of random overlap, PCC values were obtained for each spread after rotating the red and green fluorescence images through 90° relative to each other. Note that the median PCC values for experimentally observed (E) and randomized (R) fluorescence overlaps in all three plots in **(B)** are below 0.3, the lower threshold set for 'partial overlap' (see Figure 8).

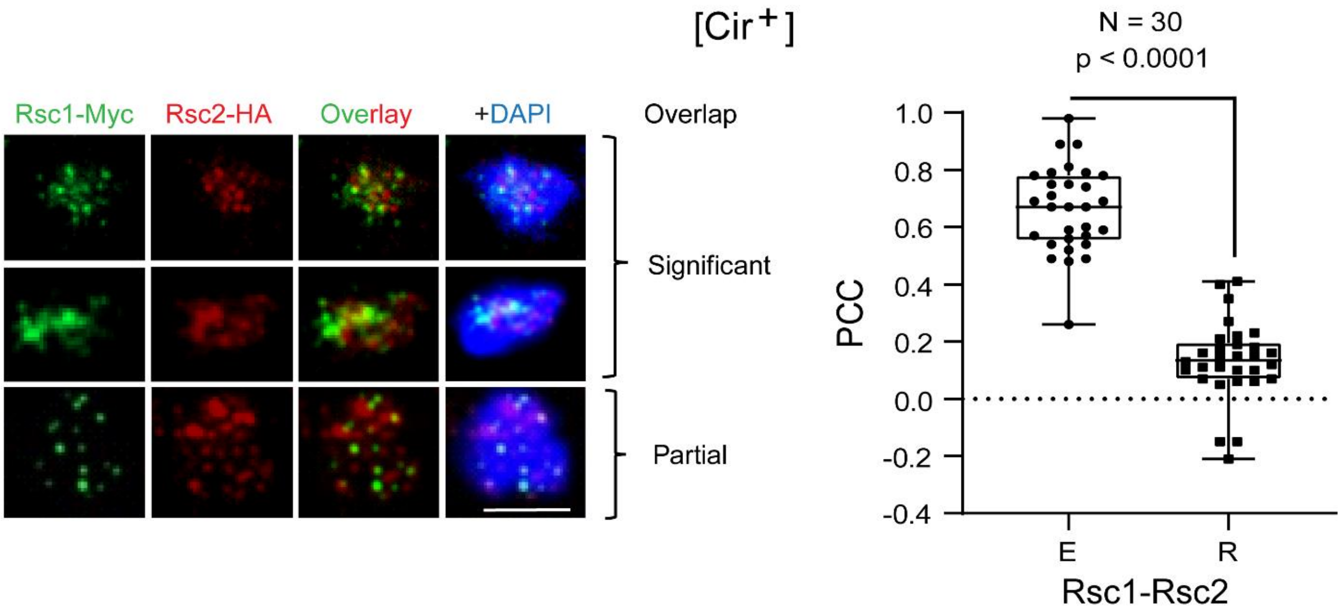

**Figure S8–2. Overlap between Rsc1 and Rsc2 in mitotic chromosome spreads.** The epitope-tagged Rsc1 and Rsc2 were visualized by immunofluorescence in [Cir$^+$] mitotic spreads. The extent of the experimentally observed Rsc1-Rsc2 fluorescence overlap (E) and the overlap following randomization (R) are plotted (see Figure 8). Bar = 5 µm.

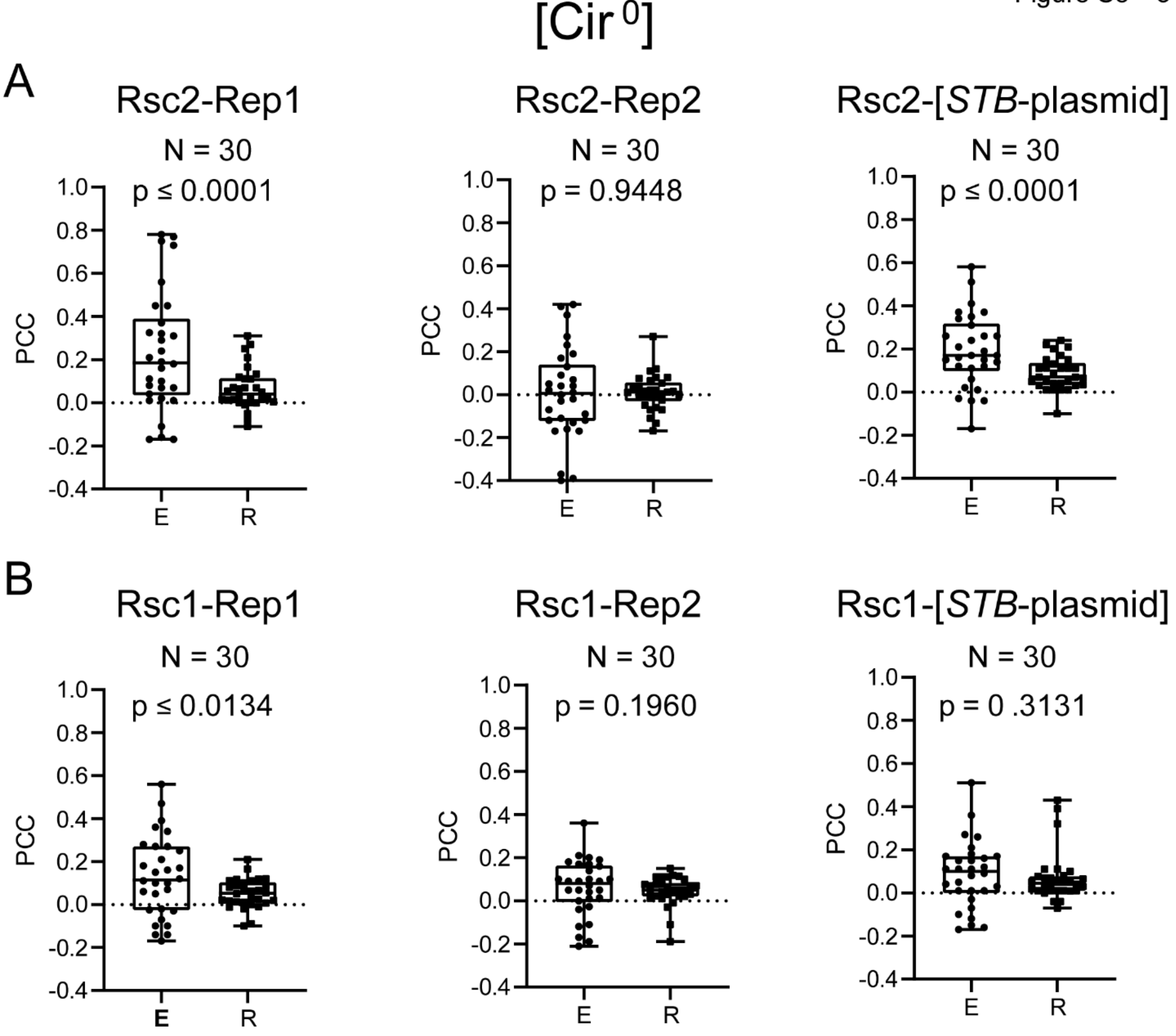

**Figure S8–3. Rsc1 or Rsc2 fluorescence overlap with Rep1 or Rep2 fluorescence in [Cir$^0$] chromosome spreads relative to fluorescence overlap by chance. (A, B)** The analysis was done as described in the legend to Figure S8–1, except that [Cir$^0$] chromosome spreads were assayed. The median PCC values for fluorescence overlap in the experimental (E) and randomized (R) samples in all the plots are < 0.3, signifying no overlap (see Figure 8).

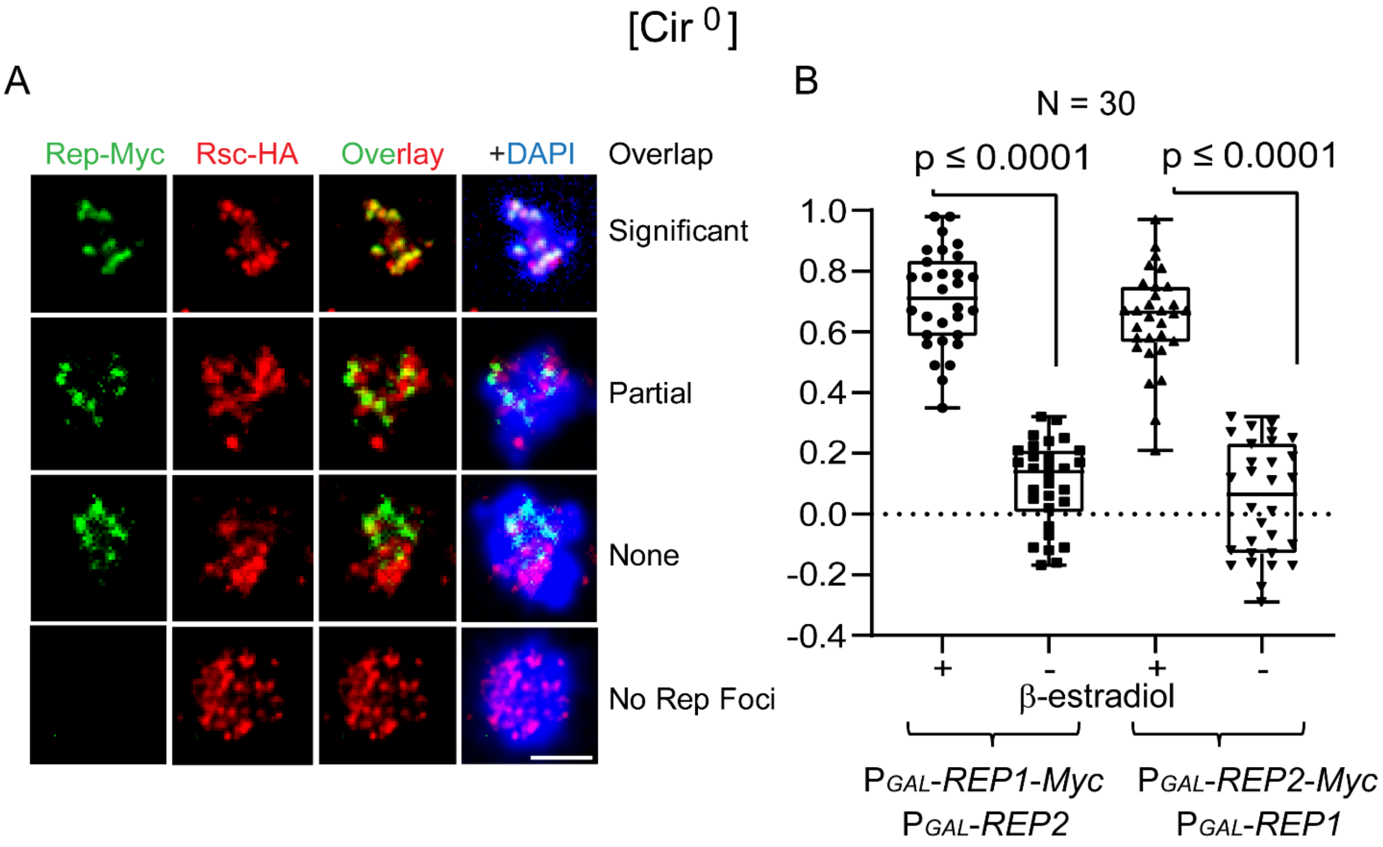

**Figure S8–4. Localization of Rep1 and Rep2 with respect to Rsc2 in meiotic chromosome spreads.** Chromosome spreads were prepared from [Cir$^{0}$] diploid cells transferred to sporulation medium for 8 hr, and were assayed by immunofluorescence microscopy. The native *RSC2* locus was modified to express Rsc2-HA. Rep1-Myc or Rep2-Myc, expressed under *GAL* promoter control from an integrated cassette, was complemented by its untagged Rep partner expressed by a *CEN*-plasmid from the *GAL* promoter. A β-estradiol inducible activator system (Carlile and Amon, 2008) was used to control the *GAL* promoter. Bar 5 μm.

Figure S9

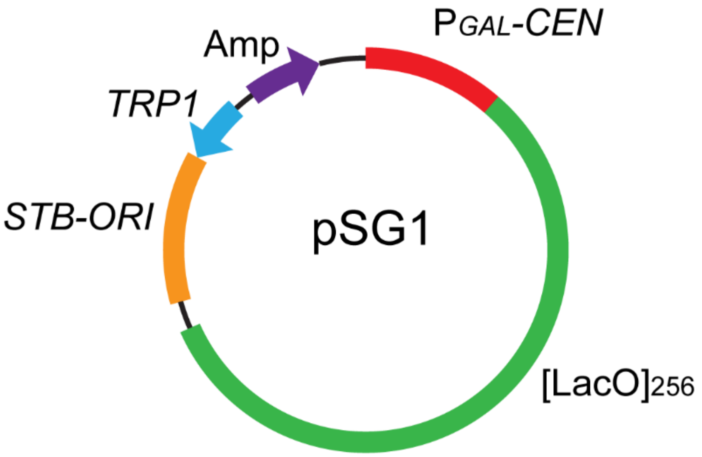

| | Host | Carbon source |
|---|---|---|
| pSG1-*CEN* | [Cir$^0$] | Glucose |
| pSG1-*ARS* | [Cir$^0$] | Galactose |
| pSG1-*STB* | [Cir$^0$] P*GAL*-*REP1* P*GAL*-*REP2* | Galactose |

**Figure S9. A nearly single-copy yeast plasmid whose partitioning capability can be modified by manipulating the host strain and the carbon source.** The features of the pSG1 plasmid are schematically diagrammed. In addition to the *TRP1* marker for selection in yeast and $[LacO]_{256}$, the plasmid carries the *ORI*-*STB* sequence from the 2-micron plasmid and a *CEN* sequence whose function is regulated by the *GAL* promoter (Ghosh et al., 2007). The plasmid behaves as pSG1-*CEN* in a [Cir$^0$] host grown in glucose (Rep1-Rep2 proteins absent; *CEN* active) and as pSG1-*ARS* when this strain is shifted to galactose (*CEN* inactive). In a [Cir$^0$] host expressing Rep1 and Rep2 under *GAL* promoter control, the plasmid is pSG1-*STB* in the presence of galactose.
